# An improved vector for baculovirus-mediated protein production in mammalian cells

**DOI:** 10.1101/2021.10.18.464913

**Authors:** Wenjun Guo, Mengmeng Wang, Lei Chen

## Abstract

BacMam system utilizes baculovirus to deliver exogenous genes into mammalian cells and is extensively used for recombinant production of eukaryotic proteins. Here, we described an improved BacMam vector (pBMCL1, Addgene#178203) which allows convenient tracing of virus production, provides higher infection efficiency towards mammalian cells, minimizes unwanted transcription of toxic genes in insect cells, and provides the capability for co-expression of multiple proteins via a single virus. We demonstrate the successful application of the pBMCL1 vector for the expression of not only the human TRPC3 channel but also the heteromeric KATP channel.

## 1. Introduction

Baculoviruses are double-stranded DNA viruses that can infect and replicate in insect populations. *Autographa californica* multiple nuclear polyhedrosis virus (AcMNPV) is one of the most common isolates of baculovirus. A modified version of AcMNPV, of which the polyhedrin gene is replaced by a target gene, is widely used for over-expression of recombinant protein in insect cells. Since it was found that the modified AcMNPV could mediate gene-transfer in mammalian cells ^1-3^, baculovirus is more and more extensively used as a vehicle for recombinant protein production in mammalian cells and this procedure is called BacMam system. BacMam system has several advantages over other eukaryotic expression systems, including insect cells or yeast expression systems. BacMam system allows exogenous gene transcription from several available strong promoters such as CMV in mammalian cells. Moreover, the BacMam system provides genuine mammalian cell environment for the posttranslational modifications of target proteins, and the glycosylation pattern of membrane protein or secreted protein can be modified using available cell line such as GnT1^-4^. Furthermore, BacMam system often provides high level of membrane protein production compared to other expression systems, allowing the access to milligram-level purified sample of previously intractable membrane protein targeted for structural or biochemical studies ^5^.

On the membrane envelope of baculovirus, there are several copies of GP64 surface protein which might bind to the heparin sulfate proteoglycans on the plasma membrane of mammalian cells ^6^. The virus then enters the cell through clathrin-mediated endocytosis process ^7^. Many exogenous membrane proteins can be functionally displayed on the baculovirus envelope during virus assembly and budding. It is reported that the vesicular stomatitis virus glycoprotein G (VSV-G) protein displayed on baculovirus envelope can enhance the infectivity of baculovirus towards mammalian cells ^8^ and a short fragment of VSV-G (VSV-GED) is sufficient to facilitate baculovirus-mediated gene delivery in vertebrate cells ^9,10^. Moreover, the large physical size of baculovirus provides high capacity for packing exogenous DNA, allowing delivering multiple genes in one single virus. However, the current available vectors did not fully exploit these advantages of BacMam system. Here, we describe the design and application of an improved vector for BacMam system.

## 2. Materials and methods

### 2.1. Cell lines and reagents

FreeStyle 293F cells (Thermo Fisher Scientific, R79007), Sf9 cells in Sf-900 III SFM (Thermo Fisher Scientific, 12659017), FreeStyle 293 Expression Medium (Thermo Fisher Scientific, 12338018), Sf-900 III SFM medium (Thermo Fisher Scientific, 12658019), SIM SF Expression Medium, (Sino Biological Inc, MSF1), SMM 293-TI (Sino Biological Inc., M293TI), Fetal bovine serum (Vistech), PEI (Polyscience, 687600), Lauryl Maltose Neopentyl Glycol, LMNG (Antrance, NG310), Cholesteryl Hemisuccinate Tris Salt, CHS (Antrance, CH210), digitonin (Biosynth)

### 2.2. Construction of pBMCL1 vector

We synthesized two DNA sequences that contain the open reading frame of (GP64 signal peptide)-(VSV-GED)-(CreiLOV) fusion protein and (LINK1)-(CMV promoter)-(multiple cloning site, MCS)-(Woodchuck Hepatitis Virus Posttranscriptional Regulatory Element, WPRE)-(SV40 polyA signal)-(LINK2) expression cassette. We inserted (GP64 signal peptide)-(VSV-GED)-(CreiLOV) into pFastBac1 vector and replaced its SV40 polyA signal with HSVtkPolyA signal to generate a vector that displays VSV-GED protein on insect cell membrane and baculovirus envelope. Then we insert the expression cassette of (LINK1)-(CMV promoter)-(MCS)-(WPRE)-(SV40 polyA signal)-(LINK2) into the vector to the opposite direction of the pH promoter to minimize mammalian gene expression in insect cells. We name this vector pBMCL1 (Figure 1).

**Figure 1.**
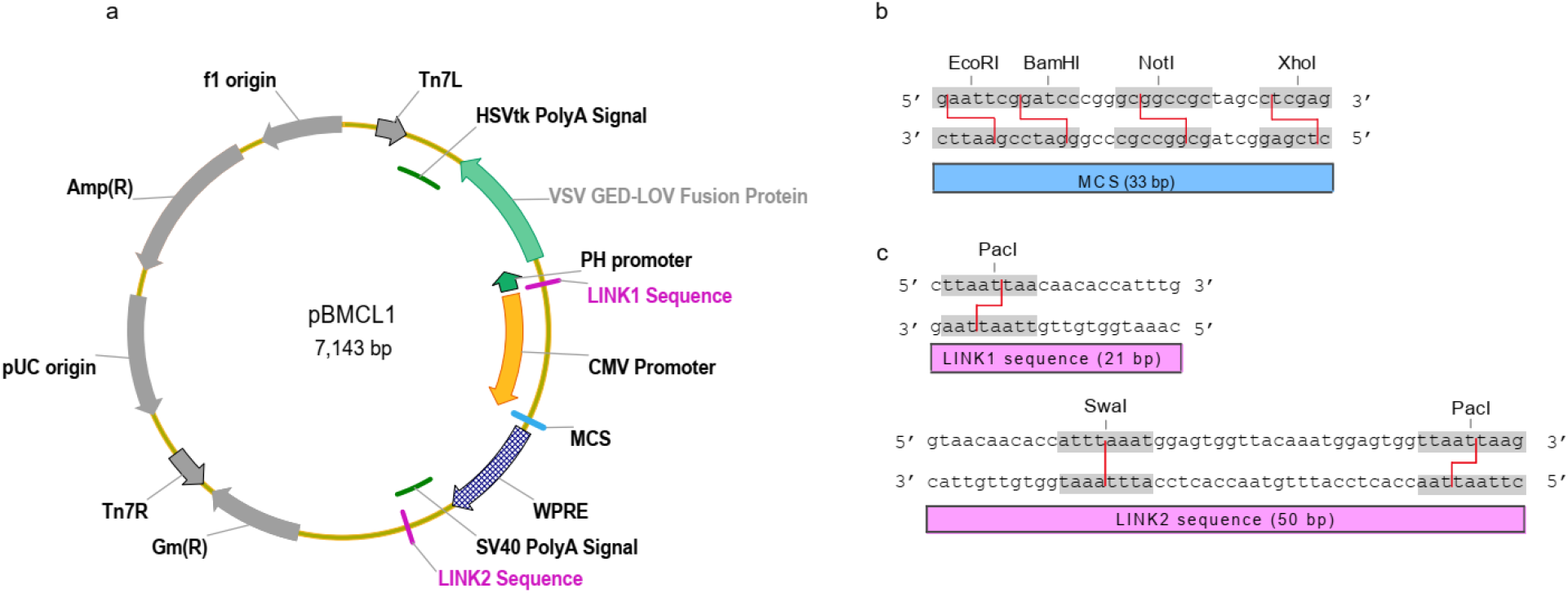
The design of pBMCL1 vector. **a**, Map of pBMCL1 vector. Elements that are essential for function are shown on the vector. **b**, The sequence of MCS and restriction enzyme recognition sites. **c**, LINK1 and LINK2 sequences and their restriction enzyme recognition sites by SwaI and PacI. These sequences are used for merging multiple expression cassettes into one vector by ligation-independent cloning.

### 2.3. Generation of expression constructs

The cDNA of human TRPC3 (hTRPC3) with N-terminal GFP was cloned into the MCS of pBMCL1 vector for protein expression using BacMam system. The cDNA of SUR1 of *Mesocricetus auratus* (XP_005084737) ^11^ and the cDNA of Kir6.2 with C-terminal GFP tag from *Mus musculus* ^12^ were cloned into the MCS of pBMCL1 BacMam expression vector to generate pBMCL1-Kir6.2-CGFP and pBMCL1-SUR1 vector. Then these two vectors were merged into one vector by the LINK sequences using ligation-independent cloning method ^13^ to generate the pBMCL1-K_ATP_ vector.

### 2.4. Generation of baculoviruses and infection of mammalian cells

pBMCL1 vectors containing target proteins were transformed into DH10Bac *Escherichia coli* cells. Recombinant baculoviruses were generated using the Bac-to-Bac Baculovirus Expression system (Invitrogen). Sf9 cells were cultured in Sf-900 III SFM medium (Gibco) at 27°C and seeded in 12-well plate. After transfection for 7 days, the fluorescence of CreiLOV was detected and the supernatant was collected. Higher titer P2 virus was amplified using sf9 cells cultured in SIM SF medium. At the time of cell mortality over 20%, the supernatant that containing baculovirus was collected by centrifugation and used to infect FreeStyle 293F cells.

### 2.5. Fluorescence-detection size-exclusion (FSEC)

Cells that expressed hTRPC3 channel were solubilized in TBS (20 mM Tris pH 8.0 at 4°C, 150 mM NaCl) with 10 mM MNG, 0.1% CHS and protease inhibitors including 1 μg/ml aprotinin, 1 μg/ml leupeptin, and 1 μg/ml pepstatin on ice for 30 min. Cell lysates were removed by centrifugation at 40,000 rpm for 30 min and supernatants were loaded onto Superose 6 Increase (GE Healthcare) for FSEC analysis.

Cells that expressed K_ATP_ channels were harvested and solubilized in TBS with 1% digitonin and protease inhibitors including 1 μg/ml aprotinin, 1 μg/ml leupeptin, and 1 μg/ml pepstatin on ice for 30 min. Cell lysates were removed by centrifugation at 40,000 rpm for 30 min and supernatant was loaded onto Superose 6 Increase for FSEC analysis.

### 2.6. Microscopy imaging

Imaging experiment was performed on sf9 cells 7 days post transfection of bacmid using EVOS FL imaging system. FreeStyle 293F cells (cultured in Freestyle 293 medium) were firstly plated in 35 mm dish without shaking for 30 min, then baculoviruses were added into the medium. 48 h post infection, GFP signal was detected. Green fluorescence images were obtained using Axio Image M2 microscope.

### 2.7. Electrophysiology

For patch clamp recording, pBMCL1-K_ATP_ plasmids were transfected into FreeStyle 293F cells (cultured in FreeStyle 293 medium) at density of 1×10^6^ cells/ml using PEI and cells were cultured for 32 h before recording. Patch electrodes were pulled from a horizontal microelectrode puller (P-1000, Sutter Instrument Co, USA) to a tip resistance of 2-3 MΩ. Currents were recorded using inside-out mode at +60 mV using an Axopatch 200B amplifier (Axon Instruments, USA). We used KINT solution which contained (mM):140 KCl, 1EGTA, 10 HEPES (pH 7.4, KOH) in both pipette and bath. Signals were acquired at 20 kHz and low-pass filtered at 5 kHz. Data were further analyzed with pClampFit 9.0 software and filtered at 1 kHz.

## 3. Results and discussion

### 3.1 Design of the pBMCL1 vector

We adopted the popular Bac-to-Bac system which utilizes the site-specific Tn7-mediated transposition for bacmid generation. We modified the established BacMam expression cassette ^5^ to minimize linker lengths between each elements and to remove common restriction enzyme cutting sites out of the MCS. To confer the capability to express multiple genes simultaneously, we appended LINK1 and LINK2 sequences at 5’ and 3’ of the expression cassette respectively. LINK sequences allow the unidirectional fusion of unlimited number of expression cassettes using a ligation-independent and PCR-independent cloning method ^13^. LINK sequences are found to be functional in BacMam system based on our previous successful application on the expression of DUOX1-DUOXA1 complex ^14^. The resulting expression cassette contains a 5’ LINK1 sequence, a CMV promoter, an MCS, a WPRE, a SV40 polyA signal and a 3’ LINK2 sequence.

The successful application of BacMam system highly relies on the titer of baculovirus produced from insect cells. The strong expression of some proteins, such as certain ion channels, in insect cells might be toxic and lead to low virus-titer. Therefore, we intentionally insert the aforementioned expression cassette in reverse direction to the polyhedrin promoter of pFastbac1 vector to prevent the high level transcription of target gene in insect cells. In order to trace the successful transfection of bacmid and production of baculovirus, we utilize the polyhedrin promoter to drive the expression of a fluorescent reporter, VSV-GED-LOV fusion protein in insect cells. A GP64 signal peptide directs the fusion protein to the plasma membrane of insect cell and virus envelope. VSV-GED fragment is used to enhance the infectivity of baculovirus ^9,10^.

CreiLOV is a small, thermostable, photostable, and fast-maturing monomeric flavin-based green fluorescent protein ^15^. The commonly available green channel on fluorescent microscope for GFP detection can be used to visualize the bright CreiLOV signals. If the insect cells are successfully transfected with bacmids and are producing baculovirus, they would show green signals generated from CreiLOV which is under the control of polyhedrin promoter (Figure 2).

**Figure 2.**
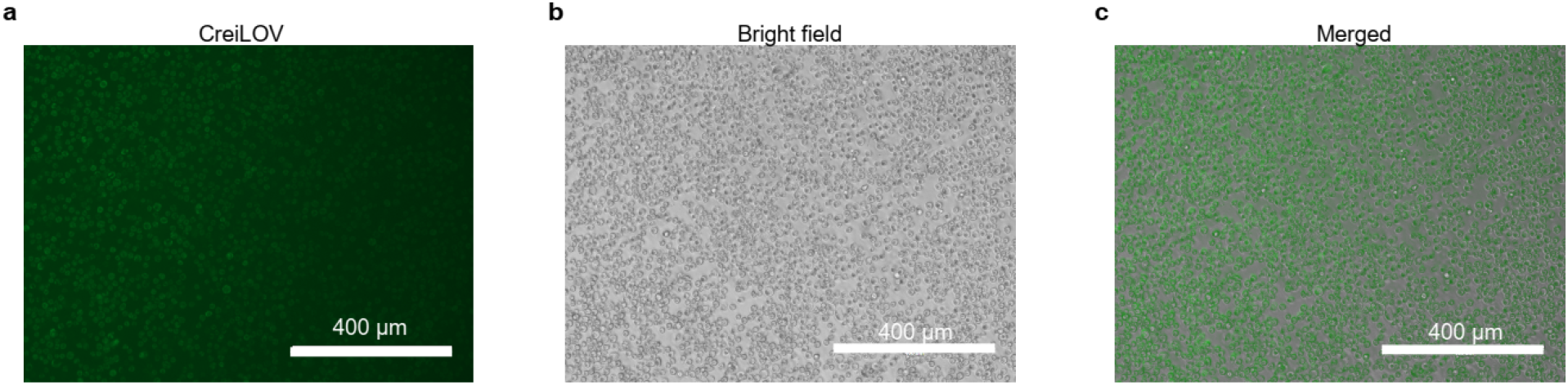
Successful baculovirus generation indicated by green fluorescence signal in sf9 cells. **a**, CreiLOV signal in sf9 cells after transfection for 7 days detected by a fluorescent microscope using green channel. **b**, Bright filed image of transfected sf9 cells. **c**, The merged image of green signal and bright field signal.

### 3.2 Using pBMCL1 vector to express human TRPC3 channel in mammalian cells

To test the function of pBMCL1 vector, we used it to express human TRPC3 (hTRPC3) channel in FreeStyle 293F cells. Full length hTRPC3 is a canonical TRP channel with molecular weight around 90 kDa. Functional hTRPC3 is a homo-tetrameric channel. We cloned the full length hTRPC3 with N-terminal GFP tag into the pBMCL1 vector. After transfection of the bacmid into sf9 insect cells for 7 days, we observed green fluorescence signal using GFP channel (Figure 3a) and P1 viruses were harvested and further amplified to obtain the P2 virus. P2 viruses were used for overexpression of hTRPC3 in FreeStyle 293F cells. 60 h post infection, we observed GFP signal from the majority of FreeStyle 293F cells (Figure 3b). We also found the band of NGFP-hTRPC3 protein at correct molecular weight on SDS-PAGE using in-gel fluorescence. (Figure 3c).

**Figure 3.**
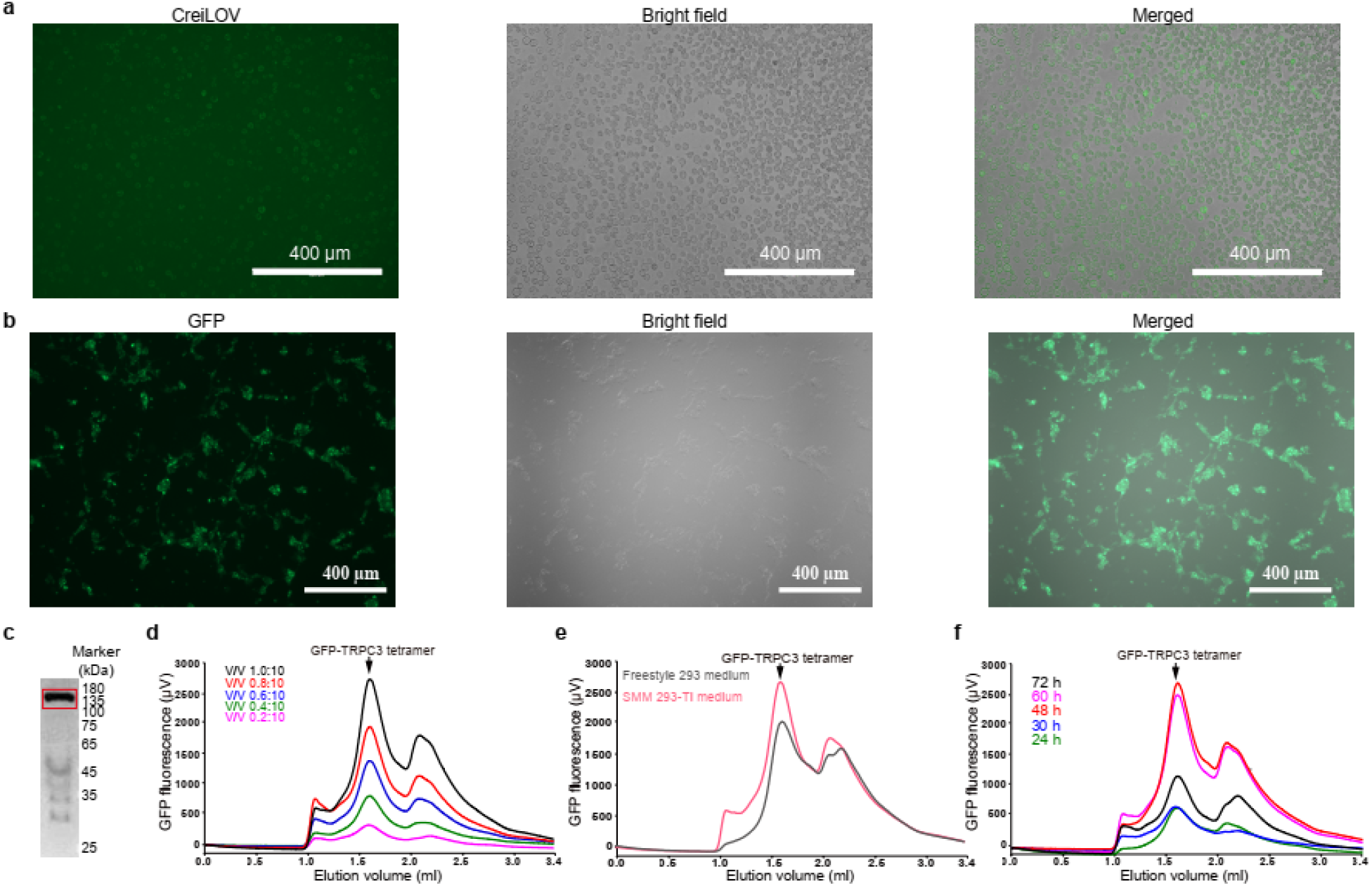
Expression of hTRPC3 channel via BacMam system using pBMCL1 vector. **a**, CreiLOV signal of sf9 cell 7 days post transfection of bacmid. **b**, GFP signal of FreeStyle 293F cells 48 h post infection by BacMam virus with virus-cell ratio 1:10 (v/v). **c**, In-gel fluorescence of NGFP-hTRPC3 protein. Band of NGFP-hTRPC3 protein is boxed in red. **d**, FSEC profiles of NGFP-hTRPC3 tetramer produced from FressStyle 293F cells infected with different virus to cell ratio (v/v). **e**, FSEC profiles of NGFP-hTRPC3 tetramer produced from FressStyle 293F cells that were cultured in different medium. **f**, FSEC profiles of NGFP-hTRPC3 tetramer produced from FressStyle 293F cells expressed for various time.

To optimize the protein expression condition, we screened different virus-cell ratio, expression duration and expression medium. We found that at the volume ratio of 1:10 (virus: cell), the expression level was highest (Figure 3d). Moreover, the expression level of hTRPC3 peaked at 48 h post-infection but decreased thereafter (Figure 3e). Furthermore, we found 293-TI medium supported higher level of hTRPC3 expression than the FreeStyle 293 medium under conditions tested (Figure 3f).

### 3.3 Using pBMCL1 vector to express pancreatic K_ATP_ channel in mammalian cells

To explore the capability of co-expressing more than one protein using a single pBMCL1 vector, we expressed the hetero-octameric pancreatic K_ATP_ channel in FreeStyle 293F cells. K_ATP_ channel is a potassium channel, the activity of which is sensitive to intracellular ATP/ADP ratio. Pancreatic K_ATP_ channel in β cells couples blood glucose level to insulin secretion and is a drug target for diabetes. Pancreatic K_ATP_ channel is assembled from four Kir6.2 subunits and four SUR1 subunits. Kir6.2 subunit is the ion channel pore with molecular weight 43.5 kDa. SUR1 subunit is the regulatory subunit with molecular weight 177 kDa. To trace the expression of K_ATP_ channel, we fused mouse Kir6.2 with a C-terminal GFP tag and cloned it into pBMCL1 vector to generate pBMCL1-Kir6.2-CGFP. We cloned non-tagged SUR1 subunit into pBMCL1 to generate pBMCL1-SUR1 vector. The expression cassette of pBMCL1-SUR1 was inserted into pBMCL1-Kir6.2-CGFP to obtain the pBMCL1-K_ATP_ vector. Transient transfection of pBMCL1-K_ATP_ into FreeStyle 293F cells allowed the recording of robust ATP-sensitive potassium currents in inside-out mode (Figure 4a). Moreover, the currents were sensitive to the insulin secretagogue repaglinide, suggesting functional K_ATP_ channels were correctly assembled and trafficked onto the plasma membrane. When solubilized, the K_ATP_ complex ran faster than Kir6.2 channel expressed alone on FSEC, indicating the hetero-octamer assembly (Figure 4b). We generated BacMam virus using pBMCL1-K_ATP_ vector for infection of FreeStyle 293F cells. We found K_ATP_ complex had robust expression 48 h post-infection (Figure 4c). We also showed the expression of both Kir6.2 and SUR1 subunits by western blot (Figure 4d). These data collectively demonstrate pBMCL1 vector is capable for the expression of eukaryotic membrane protein complex.

**Figure 4.**
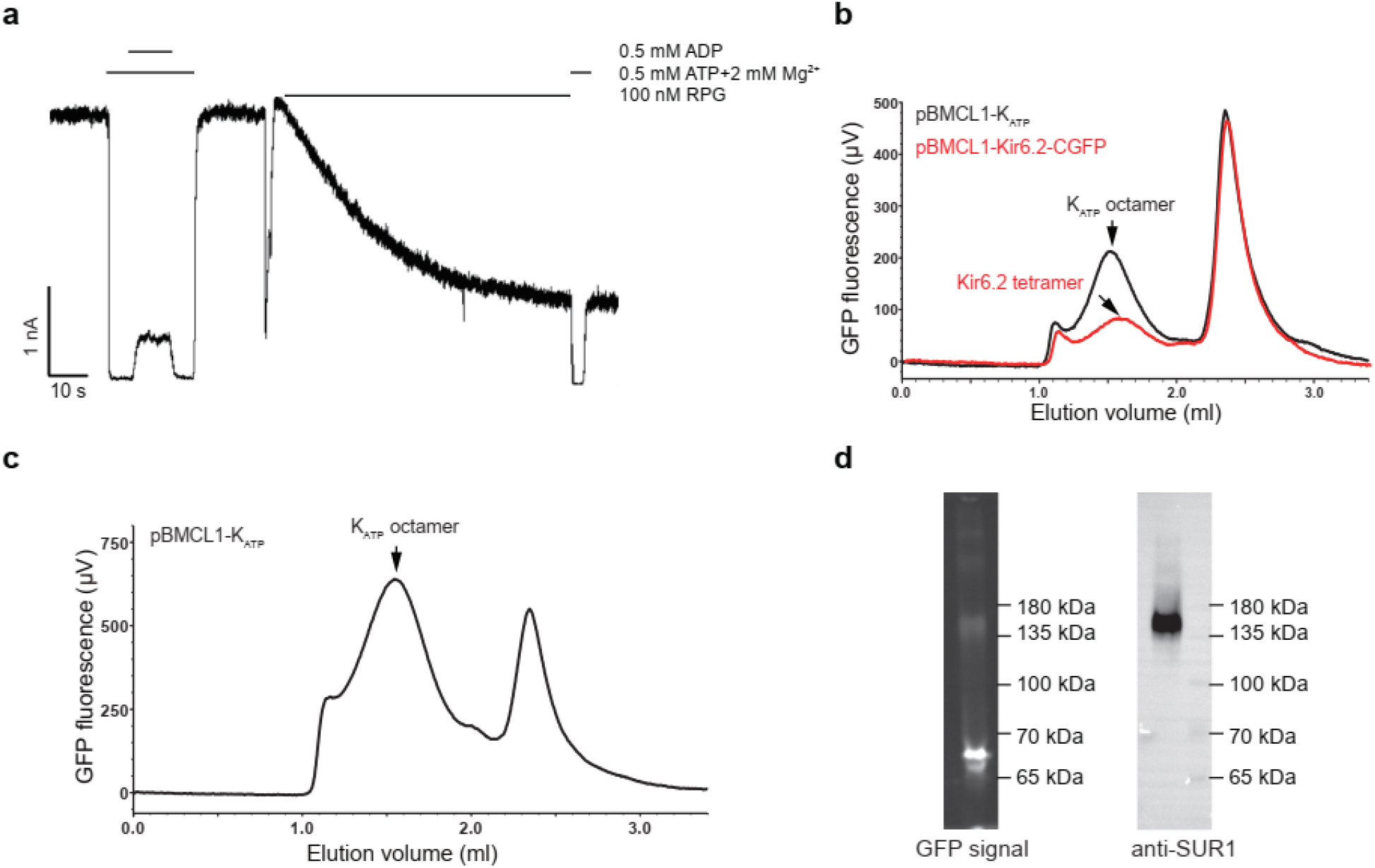
Expression of K_ATP_ channel via BacMam system using pBMCL1 vector. **a**, Recording of robust ATP-sensitive potassium currents in inside-out mode in FreeStyle 293F cells which were transfected by pBMCL1-K_ATP_ plasmid, indicating functional co-expression of both Kir6.2 and SUR1 subunits. **b**, The FSEC profile of K_ATP_ complex (using pBMCL1-K_ATP_ plasmid) and Kir6.2 channel in transfected cells. The earlier elution of K_ATP_ complex on size-exclusion chromatography suggests the successful hetero-complex formation between Kir6.2 and SUR1 subunits. **c**, The FSEC profile of K_ATP_ complex in FreeStyle 293F cells infected by pBMCL1-K_ATP_ BacMam virus. **d**, In virus-infected cells, the expression of Kir6.2 was detected by in-gel GFP fluorescence and the expression of SUR1 subunit was detected by western blot using anti-SUR1 antibody.

## 4. Conclusion

In this paper, we have described the design of pBMCL1 vector with several new features. First, this vector facilitates the tracing of successful virus generation by green fluorescence signal from LOV. Second, this vector might enhance the infectivity of BacMam virus towards mammalian cells by displaying VSV-GED fragment on baculovirus surface. Third, the expression of target protein in insect cells is minimized by placing CMV promoter in reverse direction to the polyhedrin promoter. Fourth, the incorporation of LINK sequences permits co-expression of multiple genes using BacMam system at the same time. Using this vector, we successfully expressed homo-tetrameric ion channel hTRPC3. Furthermore, we demonstrated the functional expression of hetero-octameric pancreatic K_ATP_ channel using this vector. We believe the aforementioned new features of this vector would help researchers to fully unleash the capability of BacMam system for recombinant protein expression.

## Supporting information

vector map of pBMCL1

## Credit authorship contribution statement

**Wenjun Guo:** Methodology, Validation, Formal analysis, Investigation, Writing - review & editing; **Mengmeng Wang:** Methodology, Validation, Formal analysis, Investigation, Writing - review & editing; **Lei Chen:** Conceptualization, Methodology, Validation, Formal analysis, Investigation, Writing - original draft, Writing - review & editing, Visualization, Supervision, Project administration, Funding acquisition.

## Acknowledgements

We thank Kaili Xue from Prof. Yifu Qiu lab and Peixue Xia from Prof. Ying Liu lab for kindly help of microscopy imaging. We thank Chen lab members for kindly help.

## References

1 Hofmann, C. et al. Efficient gene transfer into human hepatocytes by baculovirus vectors. Proc. Natl. Acad. Sci. U. S. A. 92, 10099–10103, doi:10.1073/pnas.92.22.10099 (1995).

2 Boyce, F. M. & Bucher, N. L. Baculovirus-mediated gene transfer into mammalian cells. Proc. Natl. Acad. Sci. U. S. A. 93, 2348–2352, doi:10.1073/pnas.93.6.2348 (1996).

3 Shoji, I. et al. Efficient gene transfer into various mammalian cells, including non-hepatic cells, by baculovirus vectors. J. Gen. Virol. 78 (Pt 10), 2657–2664, doi:10.1099/0022-1317-78-10-2657 (1997).

4 Dukkipati, A., Park, H. H., Waghray, D., Fischer, S. & Garcia, K. C. BacMam system for high-level expression of recombinant soluble and membrane glycoproteins for structural studies. Protein Expr. Purif. 62, 160–170, doi:10.1016/j.pep.2008.08.004 (2008).

5 Goehring, A. et al. Screening and large-scale expression of membrane proteins in mammalian cells for structural studies. Nat. Protoc. 9, 2574–2585, doi:10.1038/nprot.2014.173 (2014).

6 Wu, C. & Wang, S. A pH-sensitive heparin-binding sequence from Baculovirus gp64 protein is important for binding to mammalian cells but not to Sf9 insect cells. J. Virol. 86, 484–491, doi:10.1128/JVI.06357-11 (2012).

7 Long, G., Pan, X., Kormelink, R. & Vlak, J. M. Functional entry of baculovirus into insect and mammalian cells is dependent on clathrin-mediated endocytosis. J. Virol. 80, 8830–8833, doi:10.1128/JVI.00880-06 (2006).

8 Barsoum, J., Brown, R., McKee, M. & Boyce, F. M. Efficient transduction of mammalian cells by a recombinant baculovirus having the vesicular stomatitis virus G glycoprotein. Hum. Gene Ther. 8, 2011–2018, doi:10.1089/hum.1997.8.17-2011 (1997).

9 Kaikkonen, M. U. et al. Truncated vesicular stomatitis virus G protein improves baculovirus transduction efficiency in vitro and in vivo. Gene Ther. 13, 304–312, doi:10.1038/sj.gt.3302657 (2006).

10 Graves, L. P. et al. Improved Baculovirus Vectors for Transduction and Gene Expression in Human Pancreatic Islet Cells. Viruses 10, doi:10.3390/v10100574 (2018).

11 Aguilar-Bryan, L. et al. Cloning of the beta cell high-affinity sulfonylurea receptor: a regulator of insulin secretion. Science 268, 423–426 (1995).

12 Woo, S. K., Kwon, M. S., Ivanov, A., Gerzanich, V. & Simard, J. M. The sulfonylurea receptor 1 (Sur1)-transient receptor potential melastatin 4 (Trpm4) channel. J. Biol. Chem. 288, 3655–3667, doi:10.1074/jbc.M112.428219 (2013).

13 Scheich, C., Kummel, D., Soumailakakis, D., Heinemann, U. & Bussow, K. Vectors for co-expression of an unrestricted number of proteins. Nucleic Acids Res. 35, e43, doi:10.1093/nar/gkm067 (2007).

14 Wu, J. X., Liu, R., Song, K. & Chen, L. Structures of human dual oxidase 1 complex in low-calcium and high-calcium states. Nat Commun 12, 155, doi:10.1038/s41467-020-20466-9 (2021).

15 Mukherjee, A. et al. Engineering and characterization of new LOV-based fluorescent proteins from Chlamydomonas reinhardtii and Vaucheria frigida. ACS Synth Biol 4, 371–377, doi:10.1021/sb500237x (2015).

